# PsyMuKB: A *De Novo* Variant Knowledge Base Integrating Transcriptional and Translational Information to Identify Isoform-specific Mutations in Developmental Disorders

**DOI:** 10.1101/715813

**Authors:** Guan Ning Lin, Sijia Guo, Xian Tan, Weidi Wang, Wei Qian, Weichen Song, Jingru Wang, Shunying Yu, Zhen Wang, Donghong Cui, Han Wang

## Abstract

*De novo* variants (DNVs) are one of the most significant contributors to severe early-onset genetic disorders such as autism spectrum disorder, intellectual disability, and other developmental and neuropsychiatric (DNP) disorders. Currently, a plethora of DNVs have being identified through the use of next-generation sequencing and much effort has been made to understand their impact at the gene level; however, there has been little exploration of the impact at the isoform level. The brain contains a high level of alternative splicing and regulation, and exhibits a more divergent splicing program than other tissues; therefore, it is crucial to explore variants at the transcriptional regulation level to better interpret the mechanisms underlying DNP disorders. To facilitate better usage and improve the isoform-level interpretation of variants, we developed the PsyMuKB (NeuroPsychiatric Mutation Knowledge Base), a knowledge base containing a comprehensive, carefully curated list of DNVs with transcriptional and translational annotations to enable identification of isoform-specific mutations. PsyMuKB allows a flexible search of genes or variants and provides both table-based descriptions and associated visualizations, such as expression, transcript genomic structures, protein interactions, and the mutation sites mapped on the protein structures. It also provides an easy-to-use web interface, allowing users to rapidly visualize the locations and characteristics of mutations and the expression patterns of the impacted genes and isoforms. PsyMuKB thus constitutes a valuable resource for identifying tissue-specific *de novo* mutations for further functional studies of related disorders. PsyMuKB is freely accessible at http://psymukb.net.

## Introduction

In addition to inheriting half of each parent’s genome, each individual is born with a small set of novel genetic changes, referred to as *de novo* variants (DNVs), that occur during gametogenesis [1,2]. These variants, which are identified in parent-offspring trios, range in size from single-nucleotide variants (SNVs) to small insertions and deletions (indels), termed as *de novo* mutations (DNMs), and larger structural variations as *de novo* copy number variants (CNVs), and have been implicated in various human diseases [3,4]. Recently, a large number of DNVs have been discovered by whole-exome sequencing (WES) and whole-genome sequencing (WGS), and have been explored and analyzed at the gene level to assess their contributions to complex diseases [5–10]. However, isoform level information has rarely been explored for investigations.

As many as 95% of genes are subject to alternative splicing (AS), initiation, and promoter usage to produce various isoforms, increasing human transcriptomic and proteomic diversity [11,12], with approximately four to seven isoforms per gene [12,13]. An isoform is highly specific, and its expression is often restricted to certain organs, tissues, or even cell types within the same tissue [14–16]. Notably, this occurs at a high frequency in brain tissues [17,18] and regulates biological processes during neural development, including cell-fate decisions, neuronal migration, axon guidance, and synaptogenesis [19,20].

Exons are differentially used in isoforms of the same gene; therefore, it is likely that disease mutations may selectively impact only isoforms with mutation-carrying exons. Moreover, if some isoforms are not expressed in a particular developmental period or in a specific tissue, then the disease mutations affecting such isoforms may not manifest their functional impact at that period or in that tissue. Thus, correlating tissue-specific isoforms with disease mutations is an important and necessary task for refining our understanding of human diseases. Because is subject to the highest number of AS events[17,18], it is imperative to study mutations related to brain disorders at the isoform level with brain-specific expressions. However, the association between isoforms and DNMs in developmental and neuropsychiatric (DNP) disorders, such as autism (ASD), schizophrenia (SCZ), early-onset Alzheimer disorder (AD), and congenital heart disorder (CHD), has rarely been investigated on a large scale.

In this study, we present the NeuroPsychiatric Mutation Knowledge Base (PsyMuKB), a unique DNV database that we have developed. PsyMuKB serves as an integrative platform that enables exploration of the association between tissue-specific regulation and DNVs in DNP disorders (**Figure 1**). It provides a comprehensive collection of DNVs, both DNMs hitting coding and non-coding regions and *de novo* CNVs, spanning across 25 different clinical phenotypes, reported in 123 studies as May 2019, including DNP disorders such as ASD, SCZ, and early-onset AD. In addition, based on the genomic position of each mutation, transcriptional features, and the genomic structures of transcripts, we developed a novel pipeline that allows flexible filtering and exploration of isoforms that are impacted by mutation and/or brain-expressed with a user-specified selection. Finally, PsyMuKB allows the searching and browsing of genes by their IDs, symbols, or genomic coordinates, and provides detailed gene information, including descriptions and summaries, exon-intron structures of transcripts, expression of the gene and/or protein in various tissues, and protein-protein interactions (PPIs). Therefore, PsyMuKB is a comprehensive resource for exploring disease risk factors by transcriptional and translational information with associated visualizations. Herein, we describe the architectural features of PsyMuKB, including both the variants and their annotations, and a system for understanding the impact of mutations on tissue-specific isoforms, protein structures on brain-related complex disorders. It highlights novel mechanisms underlying the genetic basis of DNP disorders.

**Figure 1.**
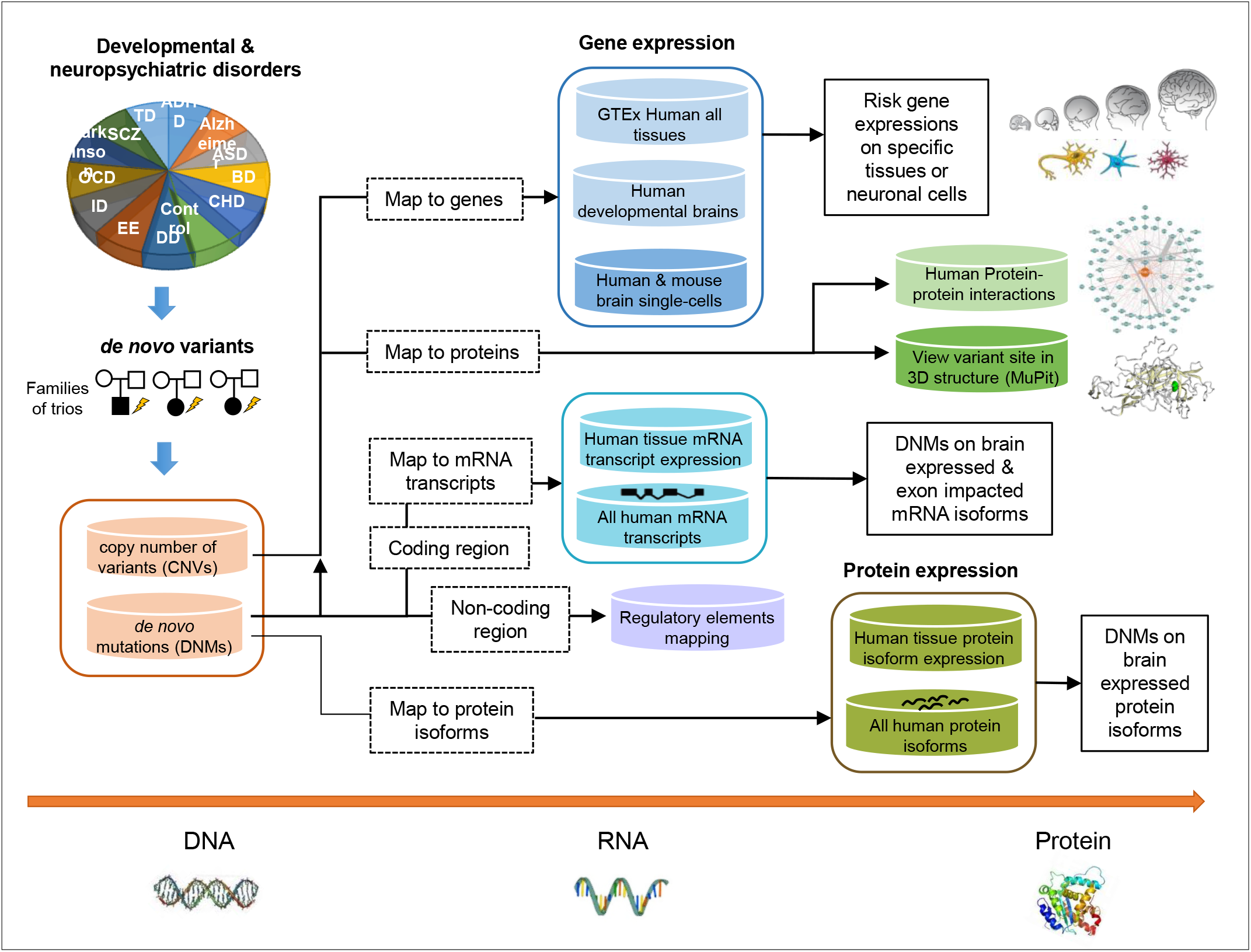
Flowchart of PsyMuKB. PsyMuKB is a multi-level DNV exploration knowledge base integrating various transcriptional and translational annotations to aid the understanding of the genetic variants from DNP disorders. Variants and genes carrying variants can be browsed, searched and filtered at gene, protein, mRNA isoform, or protein isoform levels with related annotations, such as tissue and neuronal cell-specific expression patterns, protein interactions, mutation sites mapping to regulatory elements, and protein 3D structures by muPIT [40], etc. The isoform-specific mutation filtering, using tissue-specific expression patterns and isoform exon-intron structure information, is one of key functionalities in PsyMuKB.

## Data Collection and Processing

### DNV curation

PsyMuKB catalogues two types of DNVs: (1) DNMs that include *de novo* point mutations and small indels; (2) *de novo* CNVs that involve deletions or duplications in copy numbers of specific regions of DNA. We first surveyed the literature for all published studies where human DNVs, including DNMs and CNVs, had been identified at a genome-wide scale [21]. All studies were then carefully curated to maintain essential information on each DNV, including sample identifier (if available), chromosomal locations of the reference and alternative alleles, validation status. All variants’ coordinates are shown in GRCh37 (hg19) in PsyMuKB for both DNMs and *de novo* CNVs. If source variant coordinates were not originally provided in GRCh37, the coordinates were then lifted over using the “LiftOver” from the UCSC genome browser (https://genome.ucsc.edu/cgi-bin/hgLiftOver) for annotation consistency.

The vast majority of DNM studies published and included in PsyMuKB have employed large-scale parallel sequencing using mostly WES but sometimes WGS, in conjunction with large sample sizes (hundreds to thousands of samples). These were collected from mostly from trios families, but sometimes quads families [21]. By comparing the DNA sequences obtained from affected children to those from their parents, it is possible to identify DNMs after filtering out sequencing artifacts and variant-calling errors. The variant-calling process requires a detailed bioinformatics pipeline involving the application of different thresholds to filter for various quality parameters, such as allele balance (e.g. AB between 0.3 and 0.7), allele depth (e.g. DP≥20), genotype quality (e.g. GQ≥20), mapping quality (e.g. MQ≥30), allele frequency in general population (usually <1% or 0.1% as a more stringent cutoff), etc.[5,22]. Nonetheless, all DNMs (or randomly selected subsets) are re-sequenced by other methods, usually Sanger sequencing, to check the accuracy of the findings. As a result, the average rate of DNM is estimated to be 1-3 per individual in whole exome and 60-80 per individual in the whole genome[23]. During our data collection and curation process, we ensured all the DNM data included in PsyMuKB came from discovery pipelines with reasonable quality parameters, such as those used in the 2018 study by Werling *et al*. [5]. Next, all the collected DNMs were batch-processed for systematic annotations using the ANNOVAR annotation platform[24] to include annotations, such as variant function (exonic, intronic, intergenic, UTR, etc.), exonic variant function (non-synonymous, synonymous, etc.), amino acid changes, frequency in the 1000 genome and ExAC database [25], and variant functional predictions by SIFT [26], Polyphen2 [27], GERP++ [28], and CADD [29]. Since the emphasis of many available functional annotations of variants is on coding regions, we included the DeepSea scores in the variant annotation table to help users evaluate the impact of the variants at non-coding locations. In addition, for each gene we included the Haploinsufficiency Score [30] for assessing the likelihood of the gene exhibiting haploinsufficiency and the pLI score [25] for assessing the probability of it being intolerant to loss-of-function (LoF) variants.

### Collection and processing of expression datasets

PsyMuKB includes five different datasets for expression annotations, of which four are transcriptomic data, and one is protein expression data. We selected four large-scale transcriptomic study datasets to comprehensively annotate and illustrate transcriptional expressions, including human tissue expressions from the Genotype-Tissue Expression (GTEx) consortium[31] (http://www.gtexportal.org/home/), the BrainSpan Atlas of the Developing Human Brain[32] (www.brainspan.org), and human embryonic prefrontal cortex single cell expressions [33]. Considering the majority of developmental regulation modules are preserved between human and mouse [34], we also integrated adult mouse brain single-cell expression atlas data (DropViz: http://dropviz.org/) [35], to expand the interpretive annotations of genes associated with DNVs. Gene expression levels were summarized as either Reads Per Kilobase Million (RPKM) or Transcripts Per Million (TPM) as provided by their respective sources, then we calculated and visualized all the expression levels by either original or log10-based normalized values. The BrainSpan data were plotted across six brain regions and nine developmental periods, while GTEx data were plotted by listing all human tissues in alphabetical order. All neuronal cell types were annotated by their major cell types, such as neuron, interneuron, microglia, stem cell, oligodendrocyte progenitor cell (OPC), astrocyte, etc. The human brain single-cell expressions were visualized by developmental periods and cell types, while the mouse brain single-cell expressions were visualized by brain regions and cell types. These gene expression patterns mainly aid exploration of the role of a gene in normal tissues or developmental periods, but no specific transcripts of the gene were revealed in abnormal situations. We then focused on the transcripts where DNMs were mapped to their exon locations, and where the specific location in the brain where they were expressed was recorded. To associate mutations with the brain-expressed transcripts, we mapped the genomic locations of DNMs to the exon-intron structures of each gene isoform expressed.

To associate the mutations with the protein-level annotations, we extracted the protein isoform expression data of various human tissues from ProteomicsDB (https://www.proteomicsdb.org/). Protein isoform expression data were directly extracted from ProteomicsDB with median log 10-based normalized iBAQ intensities as the expression levels. To associate the mutations with the protein isoforms expressed in the brain, we first mapped the mutation genomic locations to all the Gencode mRNA transcripts. Then, we linked Gencode mRNA transcript IDs and UniProt IDs, which were used to identify protein isoform expression data provided by ProteomicsDB. After this, we mapped the expression data to all proteins and their isoforms by UniProt IDs, and all protein expression information was plotted as histograms by different tissue type, e.g. the brain.

### Regulatory element curation and mutation mapping

Currently, functional annotations mostly emphasize mutations in coding regions. However, more than 90% of all the reported DNMs are located in non-coding regions of the genome (**Figure 2A**), which can be potentially functionally important due to the sheer size. To facilitate the usage of these variants and better explore the potential impact from the mutations hitting the non-translated genomic regions, PsyMuKB provides regulatory element annotations to help investigation of whether a non-coding mutation hits a regulatory element, potentially influencing downstream gene/isoform targets. This information is located at “Transcripts” subsection of the “Gene Information” page. There were 250,733 gene enhancer regions defined by GeneHancer[36] and 82,149 promoters defined in phase 2 of FANTOM5[37]. We have mapped curated DNMs locating at non-coding regions of the genome to all the regulatory regions and list them as part of the mutation annotations (**Figure 3**).

**Figure 2.**
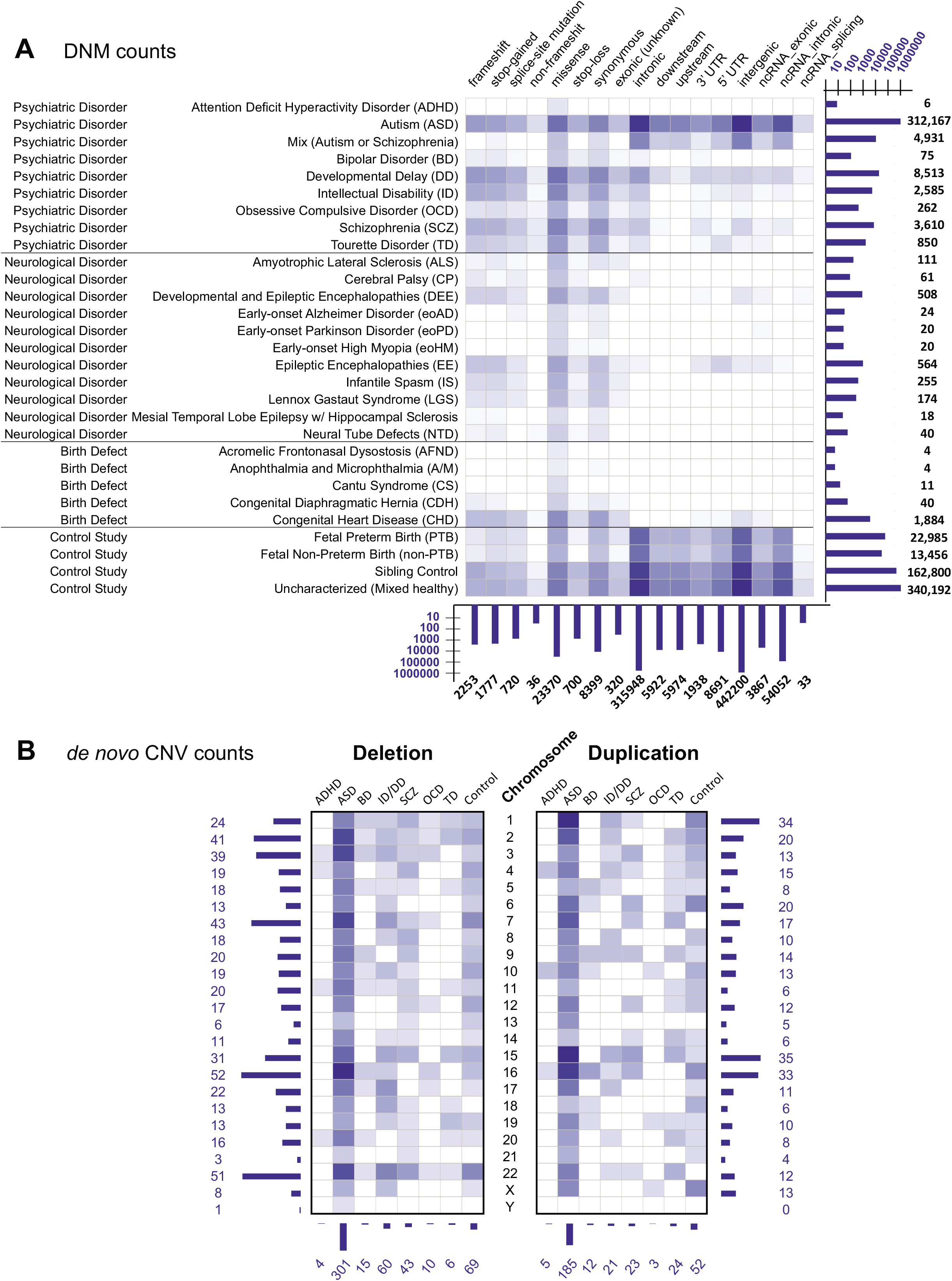
Statistics of the DNVs collected by PsyMuKB. **A.** The heatmap shows the statistics of DNMs currently in PsyMuKB. The DNMs are divided into four different clinical phenotypes (in rows): psychiatric disorder, neurological disorder, birth defect disorder and control study. The functional categories of DNMs are separated into 17 different types (in columns), from impacting coding regions (frameshift, stop-gained, spliced-site, non-frameshift, missense, stop-loss, synonymous, and exonic unknown), to noncoding regions (intronic, downstream, upstream, 5′-UTR, 3′-UTR, intergenic, and ncRNA as non-coding RNA). **B.** The heatmap shows the statistics of *de novo* CNVs currently in PsyMuKB. CNVs are collected from nine different clinical phenotypes, including ADHD, ASD, and SCZ (in columns). The figure is separated into two panels. The left represents deletion events in CNVs, and the right panel represents the duplication events in CNVs. The color in the heatmap represent the counts of each phenotype for a particular variation type, with the darker the color, the higher the count.

**Figure 3.**
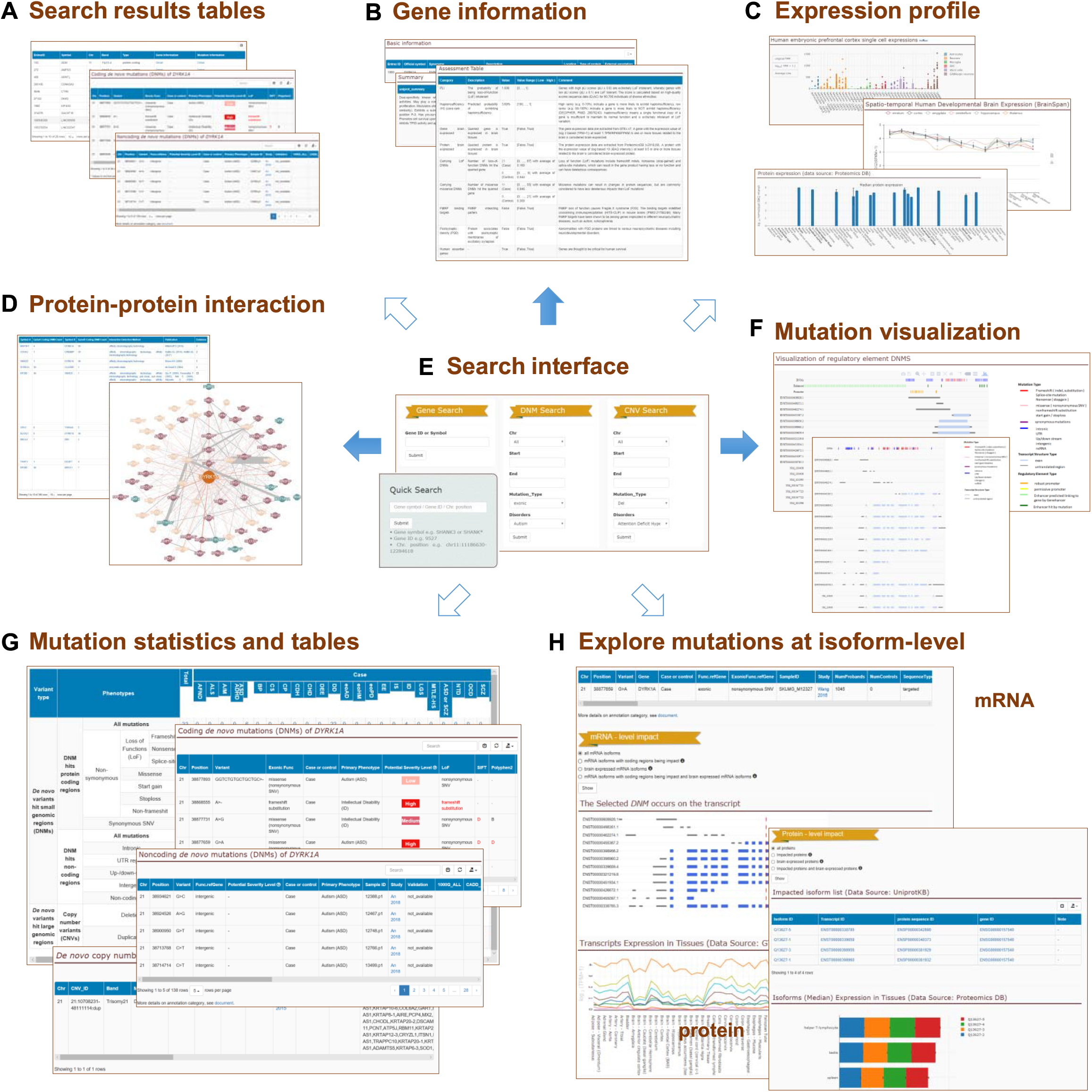
Web-interface of PsyMuKB. A snapshot of the major functionalities of PsyMuKB using neuropsychiatric-related gene *DYRK1A* as an example. **A.** The table version of the results for searching a gene/CNV/genomic location. **B.** The summary of the queried gene, such as description, synonyms, chromosomal location, summaries from RefSeq and UniProt, etc. **C.** The tissue and neuronal-specific expression profiles from various data source, such as GTEx [31], BrainSpan [32], human embryonic prefrontal cortex single cells [33], and ProteomicsDB (https://www.proteomicsdb.org/), etc. of *DYRK1A* were drawn in either histogram or curve formats to showcase the expression patterns. **D.** PPI network of *DYRK1A* was presented in both table and network-visualization formats, with the line thickness used to indicate the evidence count for a particular interaction. **E.** The main search interface of PsyMuKB. **F.** The mutation mapping in different *DYRK1A* transcripts, with regulatory elements (promoters and enhancers) also drawn to the approximated locations. **G.** The mutation statistics and result tables for *DYRK1A*. The statistics table shows the summary of all the DYRK1A variants from different mutation types in different clinical phenotypes. The DNM table details each DNM with annotations, including location, mutation changes, exonic functions, allele frequencies, deleteriousness predictions from different tools, etc. **H.** The interface for identifying the isoform-specific mutation using the tissue-specific expression of isoforms and their exon-intron structure map.

### Interaction data curation

We extracted PPI data from BioGRID [38] to construct a comprehensive map of physically interacting human proteins. After removing non-physical interactions as defined in BioGRID, we obtained 409,173 human PPIs for annotation integration, allowing users to explore the potential functional pathways involving the proteins impacted. For each interaction, we have kept the annotations, such as official symbols of both protein interactors, experiment detection method, and publication PMID.

### Database architecture

PsyMuKB has been designed as an expandable big data platform using MongoDB, a high-performance non-SQL database management system. This provides sufficient scalability and extensibility for easy and fast data integration and module expansion in future updates. All metadata in PsyMuKB are stored in the MongoDB database, while the graphical representation, such as expression profiles, mutations mapping to the transcripts, and PPI network, are mapped and drawn in real time when related data are queried. The web interface and data visualization of PsyMuKB were implemented mostly in Python scripts based on HTML5 and Cascading Style Sheets (CSS), and JavaScript (JS). The expression data visualization and regulatory element mapping were implemented using Plotly. The interaction network visualization was implemented using Cytoscape.js [39]. Illustration of the mutation site in a 3D protein structure is provided by a link to the corresponding visualization provided by the muPIT [40] interactive web server (http://mupit.icm.jhu.edu/MuPIT_Interactive/).

## Database Content and Usage

### Mutation data statistics

As of May 2019, PsyMuKB contains 877,016 DNVs in the current psymukb.v.1.5 version, covering 24 different types of brain or neuronal-related disorders and some control population studies (**Figure 2A, Data collection and processing**). A total of 876,175 of them are DNMs, including SNVs and small indels, affecting 732,879 unique sites across the genome (**Figure 2A**). About 61.5% of variants come from controls, including healthy sibling of patients (n=162,800) from various DNP disorder studies, an uncharacterized cohort study (n=340,192) [41] and a fetal sample (preterm and non-preterm) study (n=36,441) [42]. DNM variants were collected from various studies based on four major clinical phenotypes: psychiatric disorders, neurological disorders, birth defect diseases, and control studies (**Figure 2A**). In eight major developmental psychiatric disorders, the majority (93.7%) of DNMs came from ASD studies (n=312,167), followed by studies of developmental delay (DD) (n=8,513), SCZ (n=3,610) and intellectual disability (n=2,585). In neurological disorders, the majority of DNMs came from epileptic encephalopathies (EE) (n=564) and developmental and epileptic encephalopathies (DEE) (n=508). In birth-defect diseases, the majority of DNMs came from congenital heart disease (97%, n=1,884). For DNMs, half of the variants were located in intergenic regions (n=442,200), compared to only about 4.3% (n=28,259) of mutations impacting protein translation exonic regions and 38.7% located at 5′-UTR, 3′-UTR, intronic, upstream or downstream regions of the transcripts, while the remaining 6.6% of DNMs were located in non-coding RNAs (**Figure 2A**).

It has been shown that CNVs have contributed significantly to the disease etiology of psychiatric disorders [43–46]. Thus, it is vital that such variants are included in the database as well. Therefore, we have curated 841 *de novo* CNVs from reported genome-scale studies, covering eight different clinical phenotypes and affecting 369 non-overlapping genomic regions (**Figure 2B, Data collection and processing**), ranging from 1Kb to 600Mb. More than half of *de novo* CNVs (28%, n=486) are ASD CNVs, followed by control (14%), intellectual disability (9.6%), and SCZ (7.8%) CNVs. In this set of curated CNV data, 61% were deletions, and 39% were duplications. Additionally, CNVs were shown to hit most frequently at regions of chromosome 16 (10%, n=85), followed by chromosomes 22, 2, 7 and 1 (**Figure 2B**).

### Novelty of PsyMuKB

PsyMuKB does not limit its collection of variants to DNMs like three existing databases, the Developmental Brain Disorder Genes Database (DBD) [47], denovo-db [48] and NPdenovo [49]. It also provides a comprehensive list of *de novo* CNVs covering eight different clinical phenotypes (**Figure 2B, Data collection and processing**), including DNP disorders, such as ASD and SCZ. DBD (https://dbd.geisingeradmi.org/) focuses on six developmental brain disorders, ASD, intellectual disability, attention deficit hyperactivity disorder (ADHD), SCZ, bipolar disorder, and epilepsy. It also presents only a limited set of 29 genes carrying missense DNMs and a set of 465 genes with pathogenic LoF (loss-of-function, such as splice-site, stop-gain, and frame-shift) in a tiered classification based on LoF count. In addition, the current version (accessed on 2019/05/05) does not provide data visualization.

While Denovo-db (http://denovo-db.gs.washington.edu/denovo-db/) maintains a large set of DNMs (~420,000 DNMs in denovo-db.v.1.6.1) from a wide range of DNP diseases, it focuses on presenting DNMs as a variant collection with all variant data giving in a tabular format with annotations such as variant locations, frequencies in other databases and pathogenicity predictions by SIFT [26], CADD [29], etc. In its current version (accessed on 2019/05/05), it is lacking additional genetic annotations such as gene and/or protein expressions and PPIs, which would allow further interpretation. In addition, like the DBD database, it does not provide any data visualization in the current version.

The third database, NPdenovo (http://www.wzgenomics.cn/NPdenovo/index.php), covers ~97,000 DNMs in both coding and non-coding regions across a limited set of neuropsychiatric disorders, ASD, intellectual disability, SCZ, EE, and controls (accessed on 2019/05/05), whereas PsyMuKB contains ~730,000 DNMs across genomes from more than 21 different brain disorders. Both NPdenovo and PsyMuKB include additional genomic and proteomic information for risk assessment, such as gene expression and PPIs. However, PsyMuKB offers interpretations of the variants at mRNA isoform, gene, and protein isoform level in a color interactive visualization in contrast to the current version of NPdenovo only offering interpretations of gene level expression data in tabular format.

Thus, our knowledge base differs from the existing databases by assembling DNVs regardless of clinical phenotypes and variant types, with integrations with various levels of genomic and proteomic annotations. PsyMuKB is intended to be an exploratory interpretation platform of all DNVs at the level of isoforms, genes, and proteins.

### PsyMuKB website

The PsyMuKB platform consists of a database and web interface with a set of options that support the searching, filtering, visualizing and sharing of queried data (**Figure 3**). The retrieving and visualization of gene-level information in PsyMuKB is achieved in three different ways (**Figure 3E**): by ‘Gene IDs’ or ‘Gene symbols’, ‘Chromosomal regions’, and ‘Variants’. A ‘Gene IDs’ or ‘Gene symbols’ search, provided in both Basic and Advanced searches, is useful in terms of retrieving gene descriptions and summaries (**Figure 3A-B**), expression (**Figure 3C**), protein interaction (**Figure 3D**), and all reported DNVs (**Figure 3F-G**) that are associated with the gene and the supporting evidence. The ‘Chromosomal regions’ search, also provided in both Basic and Advanced option, is useful when a user is interested in retrieving all the genes and variants located within a specific region. In addition, PsyMuKB also allows the user to browse through genes in the “Browse” tab by alphabetical order of their official gene symbols. The “Browse” tab also allows the user to navigate through different developmental or neuropsychiatric disorders related gene sets. Once a gene is selected, the results are shown in the same way as through the Search option.

When a user makes a gene query, PsyMuKB takes the user to a page with a table displaying all the genes with fully and partially matched IDs or gene symbols. This table provides two clickable links: “Gene Information” and “Mutation Information”. The first links to the gene information page, which contains five different sub-sections: (1) gene information section, which has details, including descriptions and function summaries; (2) gene and protein expressions in different tissues; (3) DNVs overview; (4) transcript information for the queried gene; and (5) PPIs involving the queried gene. In “Gene information” sub-section, PsyMuKB also provides an “Assessment Table”, which includes several brain- or disease-related genetic features, such as pLI score, haploinsufficiency score rank, expressed or not-expressed in the brain, etc., to help the user better understand the relationship between the gene and diseases.

DNVs can be accessed via two different approaches: (1) through the “*De novo* variants” statistic table (**Figure 3G**) of the gene information page after searching by “Gene ID” or “Gene Symbol”, the table lists all reported variants that hit the gene of interest; 2) by specifying chromosomal regions, variants types, and/or clinical phenotypes in the advanced search to narrow the results (**Figure 3E**). The variants are grouped by the genes annotated as associating with them, such as at the exonic, UTR or intronic region of a gene, or regulatory regions of one or multiple genes, or integenic (in between) of two distant genes. Thus, if a user queries a gene, all the related variants are shown together in two tables: coding mutations and non-coding mutations. The variant tables include information about the mutation, such as location, mutation type, case or control, disease phenotype, mutation site in the protein structure, validation status, frequency in major population databases (1000 genome, ExAC, gnomAD). Importantly, PsyMuKB provides a “Potential Severity Level” assessment annotation with three severity level defined: 1) High severity: a coding variant is either a LoF mutation or predicted as pathogenic (or deleterious) by at least three of five widely used pathogenicity prediction tools (SIFT, Polyphen2, GERP++, CADD, and ClinVar); 2) Medium severity: a coding variant is predicted as pathogenic (or deleterious) by one or two of five prediction tools; 3) Low severity: all other coding variant. This mutation-level assessment, together with gene-level assessment, aids a greater understanding of queried gene and the specific mutation carried by it.

PsyMuKB also provides basic genomic information on annotated regulatory elements, such as promoters and enhancers, by visualizing their locations on mRNA transcripts of the queried gene (**Figure 3F**). Moreover, all reported DNMs are mapped and visualized on top of the exon-intron structure of the mRNA transcripts, together with their regulatory elements, which may aid elucidation of the potential roles of the regulatory elements. In addition, PsyMuKB utilizes alternatively spliced isoforms with tissue-specific expression information, together with DNM mapping on top of the isoform structures in order to provide isoform-specific mutation selections (**Figure 3H**).

PsyMuKB also provides a human protein interaction map for the queried protein (**Figure 3D**). The interaction network is constructed using both first- and second-degree interactions and interactively visualized using Cytoscape.js [39]. The first-degree interactions are defined as the interactions between all proteins and the queried protein. The second-degree interactions are defined as all the interactions between the interacting protein partners of the queried protein. The line thickness for an interaction represents the number of items of supported evidence the interaction has. We define an evidence as either a single reported publication or a single supported experiment. If the number of PPI network protein nodes of the queried protein exceeds 200, only interactions with at least two evidence items are shown in the network. Besides the visualization, we provide a PPI table, which lists all the interaction information, including experiment detection methods, reported publications and total evidence count, regardless of the amount of evidence.

### Exploring mutations at the isoform-level

One of the key features of PsyMuKB is that it allows visualization of the DNM locations at the transcript-level and identification of affected isoforms with tissue-specific expression annotations, both at mRNA and protein levels. Here, we first assessed the necessity of studying DNMs at the isoform level and explored the scale of the DNMs that satisfy the criteria above. We used all mRNA transcripts from Gencode v19 and protein isoforms from UniProtKB (version released on 2018_07), which have been integrated into PsyMuKB, and defined three types of isoforms, “longest isoform”, “brain-expressed isoform” and “not brain-expressed isoform” (**Figure 4A-B**). At the mRNA level, the “longest isoform” is the isoform with the longest coding sequence compared to all other isoforms of the same gene; the “brain-expressed isoform” is an isoform with expression of TPM≥1 in at least one brain tissue from GTEx data; and the “not brain-expressed isoform” is an isoform that is not expressed (TPM<1) in any brain tissue sample from GTEx data. At the protein level, the “longest isoform” is the isoform with the longest amino acid sequence in a protein; the “brain-expressed isoform” is an isoform with expression of iBAQ intensity ≥1 in at least one brain tissue from ProteomicsDB data; and the “not brain-expressed isoform” is an isoform that is not expressed (iBAQ intensity <1) in any brain tissue sample from ProteomicsDB data.

**Figure 4.**
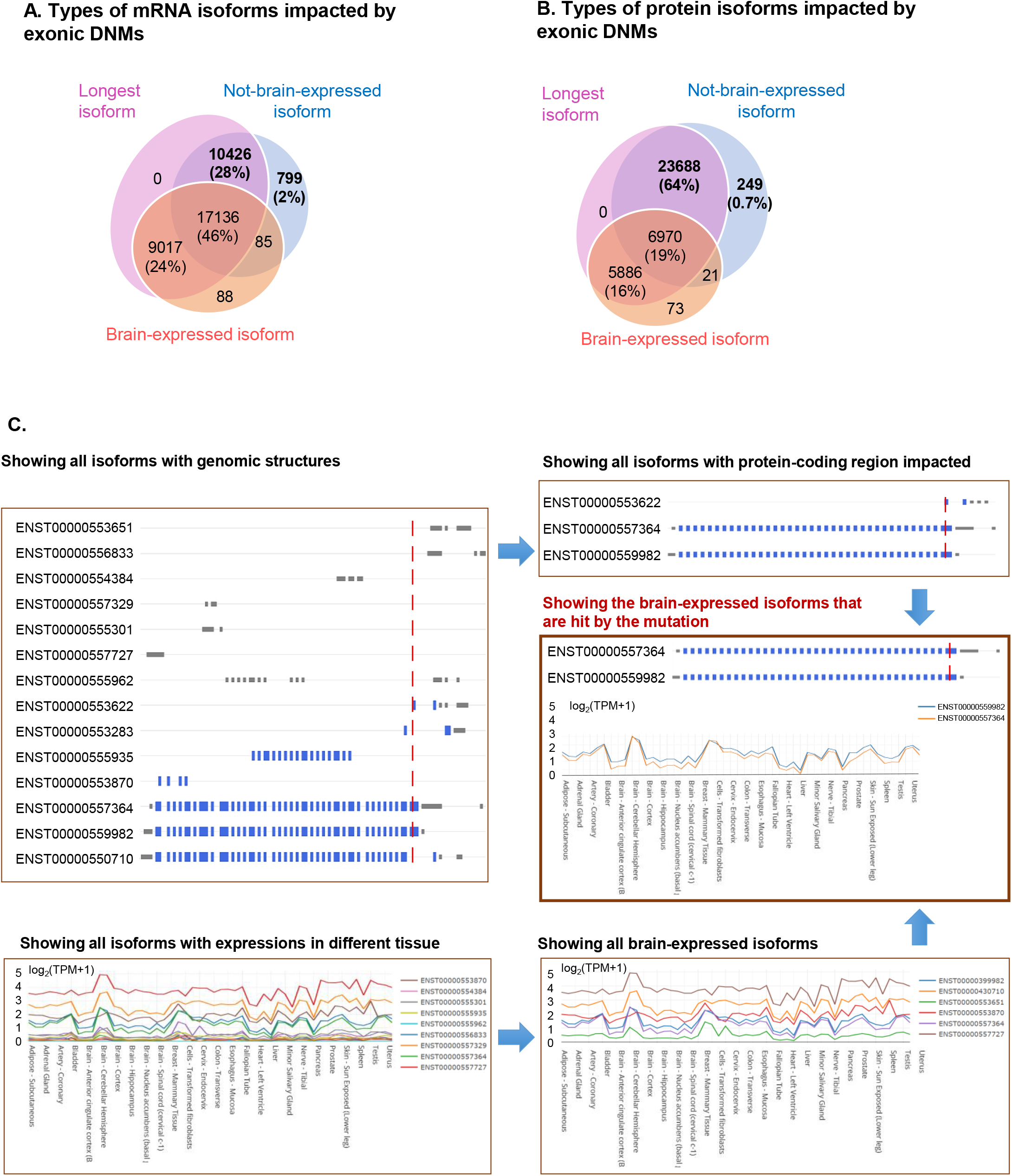
Exploration of mutations at the isoform-level. **A.** The pie chart shows the overlaps and percentages of DNMs at exonic regions hitting the three different types of mRNA isoforms, showing about 28% of DNMs only impact “not-brain-expressed” mRNA isoforms. **B.** The pie chart shows the overlaps and percentages of DNMs at exonic regions hitting the three different types of protein isoforms, showing about 64% of DNMs only impact “not-brain-expressed” protein isoforms. **C.** Example of the isoform-specific mutation filtering pipeline using a stop-gain DNM in chromosome 14 at position 21899618 with a G>C change in gene *CHD8*.

We annotated those DNMs in PsyMuKB hitting brain-expressed isoforms and identified these as “brain-expressed” mutations, as well as identifying “not-brain-expressed” mutations. Although DNMs can occur anywhere in the genome, the exome, or protein-coding region of the genome, is often investigated first when studying human disease [6,7,50]. Therefore, “not-brain-expressed” mutations may not be as interesting to researchers studying tissue-specific disease biology.

Using the “longest isoform” as the reference isoform has been a common practice in many studies and databases. Here, we ask the question whether the longest isoform strategy is still applicable for studying tissue-specific mutations. First, we looked at the exonic DNMs that impact isoforms and observed that the majority would hit the longest isoforms as expected due to the length: 97% at the mRNA-level and 99% at the protein-level (**Figure 4**). However, when checking whether most DNMs would hit at least one brain-expressed isoform, we observed that about 28% of DNMs do not hit any brain-expressed mRNA isoforms (**Figure 4A**), and as many as 64% of DNMs do not hit any brain-expressed protein isoforms (**Figure 4B**), based on the current protein isoform annotation and protein expression information from ProteomicsDB. The results show that investigation of the impact of the disease variants at the isoform-level and tissue specificity is imperative. This is a key reason for PsyMuKB to include tissue and isoform-specific expression for investigating disease-relevant mutations.

To illustrate the exploration of isoform-specific features using PsyMuKB (**Figure 3H**), we have showcased this functionality with the neuropsychiatric disease associated gene Chromodomain-helicase-DNA-binding protein 8, *CHD8* (**Figure 4C**), which has multiple alternative spliced isoforms and wide-spread expressions across many tissues. *CHD8* is believed to affect the expression of many other genes that are involved in prenatal brain development and is a strong risk factor for DNP disorders, such as ASD[51–53]. **Figure 4C** demonstrates the isoform-specific filtering process to identify suitable models for the study of mutations in *CHD8*.

## Discussion

PsyMuKB focuses on the exploration and characterization of DNV data with integrative annotations, such as isoforms, expression, protein interactions, and protein structures, and can be accessed through a user-friendly web interface (http://psymukb.net). Unlike existing databases, PsyMuKB has a ‘Mutation’ interface after a gene has been queried or browsed, which allows a unique and useful investigation of mutations with the added complexity of the alternative splicing and brain-specific expression both at mRNA and protein levels. PsyMuKB aims to be the knowledge base that takes into consideration of the isoform specificity in different tissues when exploring variants, as a specific variant could have a differential impact on alternatively spliced and regulated isoforms, both at the transcriptional and translational level; consequently, it could induce an incomplete penetrance effect. Notably, the flexibility of PsyMuKB filtering the isoforms impacted by mutation using their genomic features and expressions enhances the ability to identify the tissue-specific *de novo* events. In addition, PsyMuKB is an integrative graphical exploration platform containing a comprehensive list of DNVs, together with various types of graphical transcriptional and translational annotations, such as the detailed genomic structures of transcripts, tissue-specific expression in both genes and proteins, protein-protein interactions, and pathogenicity assessments. Thus, PsyMuKB is a comprehensive platform aiding the understanding of the impact of DNVs on developmental disorders and highlighting the novel mechanisms underlying the onset of the diseases.

## Authors’ contributions

GNL conceived and directed the PsyMuKB project. HW and DC designed the architecture. GNL, DC, and HW drafted the manuscript. XT, SG, and JW designed and implemented the different modules of the architecture. WW, WS, and WQ curated and processed all of the data for the database. ZW and SYY participated in data collection and provided clinical insights on data interpretations. All authors read, edited, and approved the final manuscript.

